# CRISPR/Cas9-mediated knockout of the ubiquitin variant Ub^KEKS^ reveals a role in regulating nucleolar structures and composition

**DOI:** 10.1101/2023.03.17.533093

**Authors:** Julie Frion, Anna Meller, Gwendoline Marbach, Dominique Lévesque, Xavier Roucou, François-Michel Boisvert

**Affiliations:** Department of Immunology and Cell Biology, Sherbrooke, QC, Canada; Department of Biochemistry and Functional Genomics, Sherbrooke, QC, Canada

**Keywords:** Ub^KEKS^, Ubiquitin variant, Data-independent Acquisition (DIA), Nucleolus, Post-Translational Modification (PTM), Apoptosis, Protein sequestration

## Abstract

Ubiquitination is a post-translational modification responsible for one of the most complex multi-layered communication and regulation system in the cell. Over the past decades, new ubiquitin variants and ubiquitin-like proteins arose to further enrich this mechanism. Among them, the recently discovered ubiquitin variant Ub^KEKS^ can specifically target several proteins and yet, functional consequences of this new modification remain unknown. The absence of Ub^KEKS^ induces accumulation of lamin A in the nucleoli, highlighting the need for deeper investigations about protein composition and functions regulation of this highly dynamic and membrane-less compartment. By using data independent acquisition mass spectrometry and microscopy, we show here that despite not impacting protein stability, Ub^KEKS^ is required to maintain normal nucleolar organization. The absence of Ub^KEKS^ increases nucleoli’s size and accentuate their circularity while disrupting dense fibrillar component and fibrillar center structures. Moreover, depletion of Ub^KEKS^ leads to distinct changes in nucleolar composition. Notably, lack of Ub^KEKS^ favors nucleolar sequestration of known apoptotic regulators such as IFI16 or p14ARF, resulting in an increase of apoptosis in Ub^KEKS^ knockout cells observed by flow cytometry and real-time cellular growth monitoring. Overall, the results presented here identifies the first cellular functions of the Ub^KEKS^ variant and lay the foundation stone to establish Ub^KEKS^ as a new universal layer of regulation in the already complex ubiquitination system.

## Introduction

Ubiquitination, which stands for the covalent binding of a highly conserved 76 amino acids ubiquitin protein (Ub), is one of the most common post-translational modification (PTM) mechanism [1][2][3]. The isopeptide linkage between Ub C-terminal domain and the targeted protein’s lysine residue, is the result of an enzymatic cascade mediated by three classes of proteins: E1 activating enzymes, E2 conjugating enzymes and E3 ligases [4][5][6][7][8][9][10]. The large number of proteins playing a role in this cascade leads to a powerful multi-layered communication system [11][12][13]. Such a system enables Ub to play a crucial role in numerous cellular mechanisms such as protein location, protein stability or even protein-protein interaction [3].

In addition to the ubiquitin genes encoding the canonical Ub protein, other variants of Ub encoded by genes that were initially annotated as pseudogenes have been reported [14]. Pseudogenes are usually described as an altered copy of genes that cannot produce a functional protein. However, Ub^KEKS^ is an example of a ubiquitin variant encoded by the pseudogene UBBP4 and acts as a fully functional PTM [14][15]. Interestingly, Ub^KEKS^ can specifically modify a wide number of proteins that differs from the canonical Ub, thanks to 4 amino acid differences: Q2K, K33E, Q49K and N60S [14]. Cells lacking expression of UBBP4 by CRISPR/Cas9 knockout display enlarged nucleoli with an accumulation of lamin A [14]. Nucleoli are highly dynamic membrane-less organelles whose morphology and size correlate with nucleolar activity and can be used to detect cellular stress [16][17][18]. Therefore, these observations suggest a role for proteins modified by Ub^KEKS^ in regulating some of the many nucleolar functions such as the production of ribosomes or proteins sequestration [14][16][19][20][21].

In correlation with the high number of mechanisms taking place in the nucleolus, many modifications, either post-transcriptional or post-translational, serve as regulators [21][22] [23][24][25][26]. Ub and Ub-like proteins have already been shown to interfere with several key mechanisms of the nucleolus. For instance, Ub has been shown to promote protein trafficking between the nucleoli and other nuclear compartment upon cellular stress [21]. Under normal conditions, Bloom syndrome helicase (BLM) eases ribosomal RNA transcription in the nucleolus by interacting with both RNA polymerase I and DNA topoisomerase I [27][28][29]. Poly-ubiquitination of BLM however allows its translocation to the nucleoplasm to engage in double strand breaks DNA repair pathway [30]. On another hand, SUMOylation of nucleolar proteins can disrupt ribosome biogenesis by targeting processing factors [26][31]. Modification of protein PELP1 by SUMO2 prevents the recruitment of key regulatory complexes to pre-60S ribosomal particles in the nucleolus, disrupting the maturation of 32S rRNA into 28S rRNA [31][32][33][34]. Finally, it is now known that several PTMs belonging to the Ub-like proteins family can build up complex regulatory interplays among ribosomal proteins [35][36]. RPL11 is a ribosomal protein that can be modified by both SUMO2 and NEDD8 [35][37]. NEDDylation of RPL11 allows its stabilization and nucleolar localization [18][36], where it plays a role in ribosome biogenesis [21]. In case of nucleolar stress, RPL11’s NEDDylation is replaced by SUMOylation [35]: RPL11 then translocates into the nucleoplasm, triggering a p53-dependent response pathway [16][18][21][35][36][38][39]. All these examples however show only a fraction of the vast regulatory system taking place in the nucleoli.

Here, we investigate the functional impacts of Ub^KEKS^ on the nucleolus using Data Independent Acquisition (DIA) mass spectrometry and microscopy. Starting from a whole cell point of view, we worked our way down to a single-cellular-compartment scale to identify potential roles for the Ub^KEKS^ variant.

## Material and Methods

### Ub^KEKS^ Knockout HeLa cells

HeLa cells which do not express Ub^KEKS^ were previously generated using the CRISPR/Cas9 method and obtained clones were validated by PCR and direct sequencing of those PCR products [14]. In this paper, Ub^KEKS^ knockout cells are referred as “HeLa 2.7” and “HeLa 4.3”, according to the chosen sgRNAs combination for invalidating UBBP4 pseudogene (Fig 1a).

**Figure 1:**
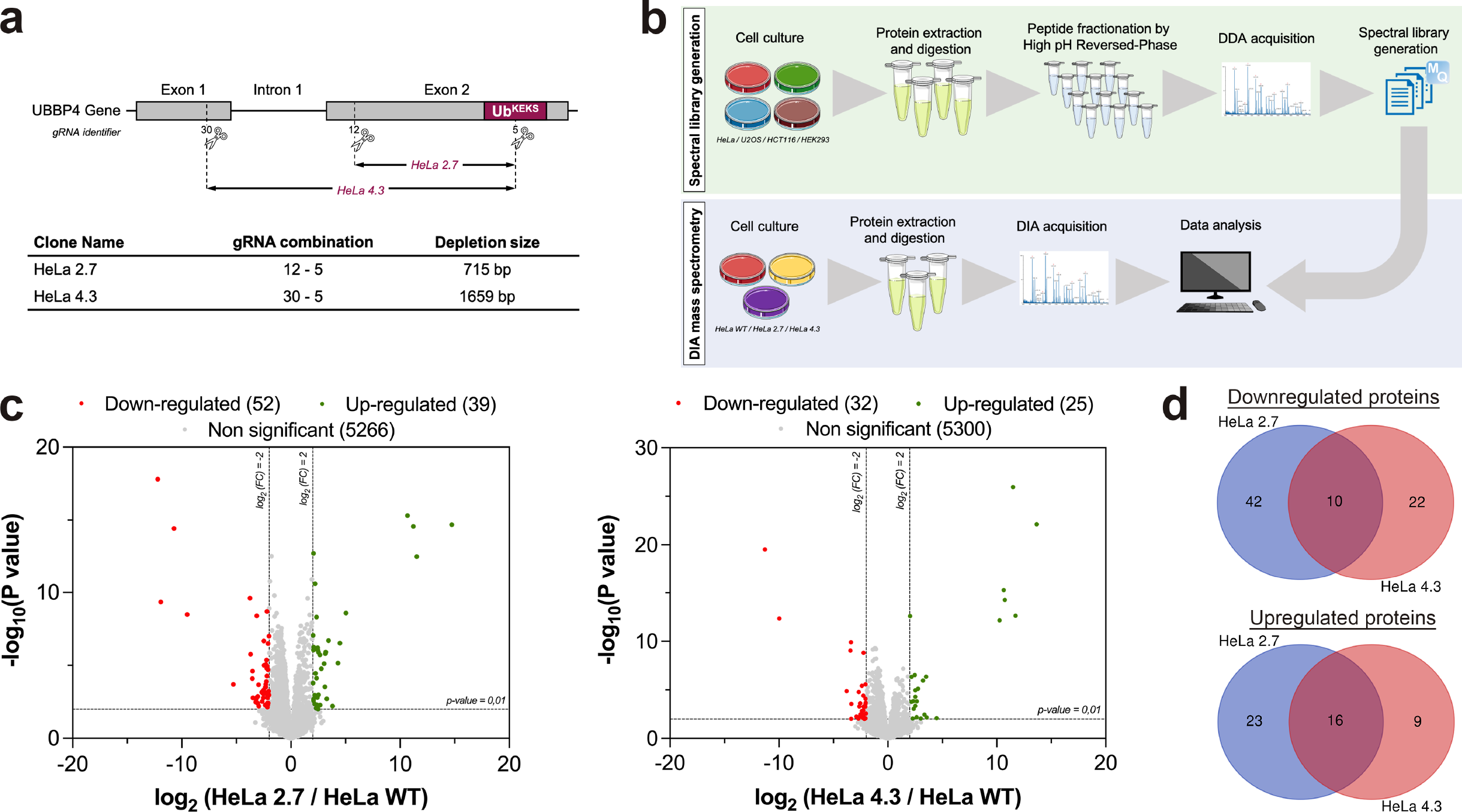
Whole cell proteome mapping of wild type and Ub^KEKS^ knockout cells. **a.** CRISPR/Cas9 strategy used to create Ub^KEKS^ knockout clones. Two distinct combinations of guide RNA were used in HeLa cells. **b.** Wild type HeLa, U2OS, HCT116 and HEK293 cells were used to create a general human spectral library for Data Independent Acquisition (DIA). Here, proteins were extracted and digested from a mix of those cell lines: data were generated using Data Dependant acquisition (DDA) and then processed with MaxQuant 2.0.3.0. DIA mass spectrometry was performed on peptides obtained from wild type and knockout HeLa cells (clones 4.3 and 2.7). The general human spectral library generated before was used to identify peptides found during the DIA acquisition. **c.** Volcano plots showing proteins identified in both wild-type and knockout HeLa cells. Differential abundance analysis was performed using a Limma t-test with slim pi0 calibration. Significantly downregulated proteins are indicated in red (fold change < -4; p-value <0.01) whereas significantly upregulated proteins are shown in green (fold change > 4; p-value < 0.01) (n=3 independent experiments, each read twice by the mass spectrometer). **d.** Overlaps between significantly downregulated or upregulated proteins in Ub^KEKS^ knockout cells. In total, 10 proteins were downregulated and 16 proteins were upregulated in both HeLa 2.7 and HeLa 4.3. The overlapping proteins list was determined using an online Venn diagram tool (Supp Fig. 2c) and later used for protein-protein network and functional enrichment analysis (Supp Fig. 2d).

### Cell culture

Wild type and Ub^KEKS^ knockout HeLa cell lines were cultured as adherent cells in Dulbecco’s Modified Eagle Medium (DMEM) which was supplemented with 10% fetal bovine serum and 100U/ml Penicillin/streptomycin.

### Study of total cell extract’s proteome

For experiments using total cell extracts, exponentially growing Wild-type and knockout HeLa cells were directly lysed into an 8M urea / 10mM HEPES pH 8 buffer. Samples were sonicated on ice (Fischer Scientific Model 120 Sonic Dismembrator) and centrifuge at 16 000g for 10 minutes (4°C) to get rid of cellular debris. In parallel, exponentially growing HEK293, HeLa, U2OS and HCT116 cells were harvested, sonicated and centrifuged following the same process in order to create a general DIA spectral library for human cells. All cleaned total cell extracts were then prepared for mass spectrometry according the steps described below.

### Immunofluorescence

Wild type and Ub^KEKS^ knockout HeLa cells were seeded on glass coverslips in 24-well plates to reach 60% confluency the next day. For rescue experiments cells were transfected with 100ng HA-Ub^KEKS^ plasmid construct using JetPrime (Polyplus-transfection SA, France) according to the manufacturer’s instructions. 24h after transfection media was changed to fresh culture media and cells were further grown for 48h. Cells were fixed on ice with cold 4% PFA (paraformaldehyde) for 20 min and permeabilized with cold 0.15% Triton X-100 in 1xPBS for 5 min. Blocking was performed using 10% goat serum (GS) in cold 1xPBS for 20 min. Coverslips were incubated with nucleolin (ab136649, Abcam, 1:2000), UBF (F-9, sc-13125, Santa Cruz Biotechnology , 1:300) or fibrillarin (38F3, ab4566, Abcam, 1:1000) primary antibodies in 10% GS in 1xPBS overnight at 4°C followed by a 1h incubation with anti-mouse (A11003, Invitrogen, 1:800) secondary antibody in 10% GS in 1xPBS at room temperature. Between each step cells were washed twice with ice cold 1xPBS. After incubation with the primary antibody, cells were washed twice with 1xPBS followed by incubation with DAPI solution (1µg/µl) for 8 min in 1xPBS and washed again twice. Coverslips were mounted on microscope slides and stored at 4°C in the dark until imaging. Images were acquired on a Zeiss LSM 880 confocal microscope using a 40× 1.4NA plan Apo objective with Z-series stacks (step size 0.45µm). Stacked images were subjected to maximum intensity projection prior analysis. Image analysis was performed using Fiji (version 1.53c) software [40]. Nucleoli circularity was measured with Fiji through object identification using nucleolin immunofluorescence data. For each nucleolus, values between 0 and 1 scale were attributed, where 0 corresponds to a line and 1 to a perfect circle. Circularity values were analyzed in Graph Pad Prism version 9.0.0. (GraphPad Software, USA).

### Transmission electron microscopy

Both wild type and Ub^KEKS^ knockout HeLa cells were grown in 6-well plates to reach 90% confluency. Cells were washed with 1xPBS and first fixed in 1.5% glutaraldehyde- Na cacodylate solution (0.1M, pH 7.4) for 30 min at room temperature followed by an overnight fixation step at 4°C in 2.5% glutaraldehyde- Na cacodylate solution (0.1M, pH 7.4). Fixed cells were washed twice in 0.1M Na cacodylate solution (pH 7.4) followed by a 1h post fixation incubation in 1% OsO4 - Na cacodylate solution (0.1M, pH 7.4). Samples were washed twice in distilled H2O and stained with 1% uranyl acetate at 4°C overnight in dark. The next day samples were washed twice in dH2O and dehydrated in a sequential manner in 40-50-70-85- 95-100% ethanol. Cells were coated with EPON epoxy resin under 25 mbar vacuum twice and polymerised at 60°C for 48 h. The specimens were then detached from the plastic dish and inverted into a new dish for embedding. After immersion in EPON epoxy resin samples were polymerized at 60°C for 48 h. After embedding, thin sections were prepared with ultramicrotome and were mounted on formvar/carbon supported copper grids (400 mesh size). Sections were contrasted with 2% uranyl acetate for 10 min and lead citrate for 5 min. Images were taken with a HITACHI H7500 electron microscope (Hitachi, Japan). All reagents were purchased from Electron Microscopy Sciences (Cedarlane, Hornby, Canada).

### RNA extraction and qPCR

RNA was isolated from wild type HeLa cells and Ub^KEKS^ knockout clones 4.3 and 2.7 cells using TRIzol reagent (Invitrogen, USA) and reverse transcription was performed using ProtoScript II reverse transcriptase (New England Biolabs, USA) with oligo dT primers. Quantitative PCR (qPCR) was performed following the manufacturer’s instruction using FastStart Essential DNA Green Master (Roche Molecular Systems, Switzerland). Target cDNAs were amplified using gene specific primer pairs (45S_FW: GAACGGTGGTGTGTCGTT, 45S_RV: GCGTCTCGTCTCGTCTCACT; 28S_FW: AGAGGTAAACGGGTGGGGTC, 28S_RV: GGGGTCGGGAGGAACGG; 18S_FW: GATGGTAGTCGCCGTGCC, 18S_RV: GCCTGCTGCCTTCCTTGG; MALAT1_FW: GACGGAGGTTGAGATGAAGC, MALAT1_RV: ATTCGGGGCTCTGTAGTCCT [41]) and *Ct* values were normalised to the expression of housekeeping genes (GAPDH_FW: TGATGACATCAAGAAGGTGGTGAA, GAPDH_RV: TCCTTGGAGGCCATGTGGGCCAT ; HPRT_FW: TGTAGCCCTCTGTGTGCTCAAG, HPRT_RV: CCTGTTGACTGGTCATTACAATAGCT) and each gene was represented as 2^-ΔΔCt^ relative to the HeLa wild type sample. The reactions were run in Optical 96-well Reaction Plates using LightCycler 96 Instrument (Roche Molecular Systems, Switzerland) and results were analysed using LightCycler 96 Software version 1.1.0.1320 (Roche Molecular Systems, Switzerland). Statistical analysis was performed using Graph Pad Prism version 9.0.0. (GraphPad Software, USA).

### Isolation of nucleoli by cellular fractionation

Nucleoli were isolated from exponentially growing wild type and KO HeLa cells in 150 mm petri dish as previously described [42]. Briefly, cells were directly lysed in 3ml of pre-chilled (-20°C) 0.5M sucrose and 3mM MgCl2 solution (=solution SI). Lysates were sonicated using the same sonicator described previously and gently layered over 3ml of a 1M sucrose and 3mM MgCl2 solution (=solution SII). Samples were centrifuged at 1800g for 10min at 4°C. Pellets were resuspended into 3ml of solution SI solution and layered on top of 3ml of solution SII and centrifuged with the same parameters for an additional wash. Purity of the nucleoli samples was checked via phase-contrast microscopy (Axiovert 200M Zeiss microscope) and the size of isolated nucleoli was measured using Fiji software version 2.1.0/1.53c [40]. Statistical analysis was performed using Graph Pad Prism version 9.0.0. (GraphPad Software, USA). Final pellets (nucleoli) were resuspended in a buffer of 8M urea, 10mM HEPES pH 8 and prepared for mass spectrometry.

### Sample preparation for mass spectrometry

Once in the 8M urea, 10mM HEPES pH 8 buffer, samples were quantified by BCA assay (Thermo Fisher Scientific #23225). DTT (5mM final concentration) was added into 50µg of proteins and samples were boiled for 2 minutes. After a 30 minutes incubation at room temperature, chloroacetamide (7,5mM final concentration) was added to the mixture. Solutions were then incubated in the dark, at room temperature for 20 minutes. 50mM of NH4HCO3 was added to each tube to reduce the final concentration of urea to 2M. Peptide digestion was performed by adding 1µg of trypsin (Trypsin Gold, Mass Spectrometry Grade, Promega Corporation, WI, USA) and incubating each sample overnight at 30°C. The samples were then acidified to a final concentration of 0.2% TFA. For samples from isolated nucleoli, a fraction of the digested sample was collected, at this step, to create the DIA spectral library generation and was processed as described in the next paragraph. On the other hand, remaining samples were cleaned using ZipTips C18 column (EMD Millipore, Burlington, VT), lyophilized in speedvac and finally resuspended in formic acid 1%. Peptides were quantified using a NanoDrop spectrophotometer (Thermo Fisher Scientific, Waltham, MA) at a wavelenght of 205 nm. The peptides were then transferred into a glass vial (Thermo Fisher Scientific) and keep at −20 °C until analysis by mass spectrometry.

### DIA spectral library generation

#### Fractionation of digested samples

Peptides collected for the DIA spectral library were desalted with ZipTips C18 column (EMD Millipore, Burlington, VT), dried in speedvac and resuspended in 300µl of 0,1% TFA. Those peptides were divided into several fractions using the Pierce High pH Reversed-Phase Peptide Fractionation kit (Thermo Fisher Scientific, Waltham, MA) according to the manufacturer protocol. Briefly, each column was initially centrifuged at 5 000g for 2 minutes at room temperature to remove the liquid and pack the resin material, followed by 2 washes with 300μl of 100% ACN. The column was then conditioned with 2 washes of 0.1% TFA. Purified peptides were loaded on the column, centrifuged at 3 000g for 2 minutes at room temperature, followed by a wash with 300μl of MS-grade water. Peptides were then eluted in 8 fractions by successively loading 300μl of 8 different solutions containing 0.1% triethylamine and 5% up to 50% ACN. For each step, a centrifugation at 3 000g for 2 minutes at room temperature was performed, with a new low-binding microtube to collect the fraction. Peptides were then concentrated by speedvac at 60°C until complete drying and then resuspended in 50 μl of 1% FA buffer. Peptides were assayed using a NanoDrop spectrophotometer (Thermo Fisher Scientific, Waltham, MA) and absorbance was measured at 205 nm. The peptides were then transferred to a glass vial (Thermo Fisher Scientific) and stored at −20 °C until analysis by mass spectrometry.

#### DDA LC–MS analysis

250 ng of peptides from each fraction were injected into an HPLC (nanoElute, Bruker Daltonics) and loaded onto a trap column with a constant flow of 4 µl/min (Acclaim PepMap100 C18 column, 0.3 mm id x 5 mm, Dionex Corporation) then eluted onto an analytical C18 Column (1.9 µm beads size, 75 µm x 25 cm, PepSep). Peptides were eluted over a 2-hour gradient of ACN (5-37%) in 0.1% FA at 400 nL/min while being injected into a TimsTOF Pro ion mobility mass spectrometer equipped with a Captive Spray nano electrospray source (Bruker Daltonics). Data was acquired using data-dependent auto-MS/MS with a 100- 1700 m/z mass range, with PASEF enabled with a number of PASEF scans set at 10 (1.17 seconds duty cycle) and a dynamic exclusion of 0.4 min, m/z dependent isolation window and collision energy of 42.0 eV. The target intensity was set to 20,000, with an intensity threshold of 2,500.

#### Protein Identification by MaxQuant Analysis using TIMS DDA [43]

The raw files were analyzed using MaxQuant (version 2.0.3.0) and the Uniprot human proteome database (version from march 2020 containing 75,776 entries). The settings used for the MaxQuant analysis (with TIMS-DDA type in group-specific parameters) were: all of the raw files were assigned with the same cell type name as well as fraction set from 1 to 8; 1 miscleavage was allowed; fixed modification was carbamidomethylation on cysteine; enzyme was set as Trypsin (K/R not before P); variable modifications included in the analysis were methionine oxidation and protein N-terminal. A mass tolerance of 20 ppm was used for both precursor and fragment ions. Identification values “PSM FDR”, “Protein FDR” and “Site decoy fraction” were set to 0.05. Minimum peptide count was set to 1. Both the “Second peptides” and “Match between runs” options were also allowed. MaxQuant was run with a transfer q value of 0.3. The “peptides.txt”, “evidence.txt” and “msms.txt” files generated from this DDA analysis were subsequently used for DIA spectral library analysis. Library quality was validated by ensuring the location of identified proteins using the COMPARTMENTS resource in the Cytoscape software environment [44][45]. For the nucleoli library, functional analysis of the 250 proteins with the highest intensity was also performed through the ShinyGO online tool (version 0.76) [46].

### DIA proteomics

#### DIA LC–MS analysis

Parameters used for DIA processing were almost identical as DDA processing, except that data were acquired using diaPASEF mode. Briefly, for each single TIMS (100 ms) in diaPASEF mode, we used 1 mobility window consisting of 27 mass steps (m/z between 114 to 1414 with a mass width of 50 Da) per cycle (1.27 seconds duty cycle). These steps cover the diagonal scan line for +2 and +3 charged peptides in the m/z-ion mobility plane.

#### Protein Identification by MaxQuant Analysis with TIMS MaxDIA [47]

Raw files were analyzed using MaxQuant (version 2.0.3.0) and the Uniprot human proteome database (version of the 21^st^ march 2020; 75,776 entries). The settings used for the MaxQuant analysis were almost identical as TIMS DDA analysis, except: TIMS MaxDIA type in group-specific parameters was selected, as well as MaxQuant as library type, followed by uploading the “peptides.txt”, “evidence.txt” and “msms.txt” files generated by MaxQuant from the specific library production. Classic LFQ normalization was also performed for every sample. Following the identification and normalization, proteins positives for at least either one of the “Reverse”, “Only identified by site” or “Potential contaminant” categories were eliminated, as well as proteins identified from a single unique peptide.

### MS data analysis

Refined tables obtained from MaxQuant were loaded into the ProStaR online interface (version 1.26.1) [48] to be further analyzed. Only identified proteins with a unique peptide count equal to or higher than 2 were considered in all analysis of this article. Normalization step of ProStaR was skipped as it was already performed during the MaxQuant analysis. Partially Observed missing values were imputated with the slsa algorithm. For value missing in the entire condition, the Det quantile algorithm (Quantile = 1; Factor =0,2) was used to impute values. Differential abundance analysis was performed with a fold change and p-value cut-off adapted to each experiment. A Limma t-test (slim pi0 calibration) was applied to determined significantly downregulated or upregulated proteins in the Ub^KEKS^ knockout clones. Volcano plots were drawn in GraphPad, based on the ProStaR differential analysis results table. Venn diagrams and the overlapping significant proteins list were obtained using an online tool available at: https://bioinformatics.psb.ugent.be/webtools/Venn/. Interactions networks from significant proteins were generated using the STRING app running in the Cytoscape software environment [45][49]. Functional analysis was also performed using the ShinyGO online visualisation tool (version 0.76) with parameters adapted for each experiment [46].

### Validation of MS data by western blot

25µg of total cell extracts or of purified nucleoli extracts were run of an 4%-20% gradient acrylamide gel (Novex™ WedgeWell™, Tris-Glycine, Invitrogen #XP04200BOX) and transferred on nitrocellulose membrane via semi-dry transfer. Two MS-identified upregulated proteins in HeLa knockout cells’ nucleoli were detected using IFI16 (ABclonal #A2007, 1:400 dilution) and P14arf (Abcam #ab216602, 1:1000 dilution) antibodies. Nucleolin (Abcam #ab136649, 1:2000) and ß-tubulin (Cell Signaling #2128S, dilution 1:1000) antibodies were also used as loading control and purity control for isolated nucleoli respectively.

### Validation of MS data by immunofluorescence

Wild type and Ub^KEKS^ knockout HeLa cells were seeded on glass coverslips in 24-well plates and prepared for confocal microscopy as described in the previous paragraph. IFI16 (ABclonal #A2007, 1:500 dilution) and P14arf (Abcam #ab216602, 1:500 dilution) antibodies were used to confirm MS results and complement immunoblotting observations. Image analysis was performed using Fiji (version 1.53c) software [40].

### Apoptosis monitoring by flow cytometry

Exponentially growing wild type and knockout HeLa cells were cultured in 6 wells plates until 80-90% confluency. Supernatants were collected and centrifuged to collect dead cells and others cellular debris. Cells were washed twice with PBS 1x and harvested by trypsinization. Cells pellets were resuspended in 100µl of Annexin V binding buffer (2.5mM CaCl2, 140mM NaCl and 10mM HEPES). Then, 5µl of PE conjugated Annexin V (Biolegend #640907) and 10µl of Propidium Iodide (0.5mg/mL stock solution) were added. Each sample were incubated for 15min in the dark prior, and their volume were completed to 500µl with Annexin V binding buffer. Signals were acquired by flow cytometry (BD Fortessa cytometer, Becton Dickinson) and analyzed with the FlowJo version 10.8.1 (Becton Dickinson & company). Percentages of cells for each of the populations went under statistical analysis in GraphPad. Significance was determined by a two-way ANOVA and Dunnett’s multiple comparisons post hoc test.

### Cellular proliferation assay

5 000 wild type and Ub^KEKS^ knockout HeLa cells were seeded and cultured as adherent cells in XCELLigence RTCA E-plates (Agilent E-plate VIEW 16, #300601140). Cells were incubated at 37°C under normal conditions. Blank with only medium was performed prior to time course measurements. Cellular impedance was measured every 15 minutes for 1 week, using a XCELLigence RTCA DP Analyzer. Growth curves were then generated from obtained raw data into Graph Pad Prism version 9.0.0. (GraphPad Software, USA). A two-way ANOVA coupled with Dunnett’s multiple comparisons post hoc test was performed to assess the significance of observed differences. At the end of the experiments, cells were fixed in 1% glutaraldehyde for 5 minutes and stained with 1% crystal violet solution. Pictures were taken using phase-contrast microscopy (Axiovert 200M Zeiss microscope).

## Results

### Whole cell proteome mapping of wild type and Ub^KEKS^ knockout cells

Ub^KEKS^ modifies different proteins compared to the canonical ubiquitin’s targets [14]. To assess the impact of endogenous Ub^KEKS^ on the whole cell scale, several Ub^KEKS^ knockout HeLa clones were generated using CRISPR/Cas9. Two distinct sets of guide RNAs (gRNAs) were used to create either a deletion of 715 bp or 1659 bp in the UBBP4 gene (Fig 1a)[14]. Whole cell proteomes of wild type and knockout cells were mapped using Data Independent Acquisition (DIA) mass spectrometry (Fig 1b). In this perspective, a complete and representative human spectral library was generated and used to increase the protein identification’s quality and reproducibility between replicates (Supp Fig 1-Supp Fig 2a). Principal Component Analysis (PCA) confirmed that Ub^KEKS^ knockout cells can be easily differentiated from wild type cells on a proteomic level (Supp Fig 2b). More than 5300 proteins were successfully identified in each sample (Fig 1c). The cellular impact of Ub^KEKS^ is well balanced with similar number of proteins downregulated or upregulated in HeLa 2.7 and in HeLa 4.3 clones. This observation is in agreement with the observation that Ub^KEKS^ does not target proteins for degradation [14]. Among the 26 proteins similarly regulated between both Ub^KEKS^ knockout clones, 10 were downregulated and 16 were upregulated (Fig 1d – Supp Fig 2c). Among the 96 proteins specifically regulated for each clone, 23 and 42 were upregulated and downregulated in HeLa 2.7, respectively, and 9 and 22 were upregulated and downregulated in HeLa 4.3, respectively. A protein-protein interaction network was generated for all significantly modulated proteins (26 proteins common to both Ub^KEKS^ knockout clones and 96 proteins specific to one clone) in Ub^KEKS^ knockout cells (Supp Fig 2d). Among the 122 proteins analyzed, 55 proteins did not have any link with the others. Also, not every protein common to both HeLa 2.7 and HeLa 4.3 had a connection. Functional enrichment analysis was performed using the online tool ShinyGo (version 0.76) and no significant function or pathway could be identified, suggesting that Ub^KEKS^ does not plays a role in protein stability or protein degradation.

**Figure 2:**
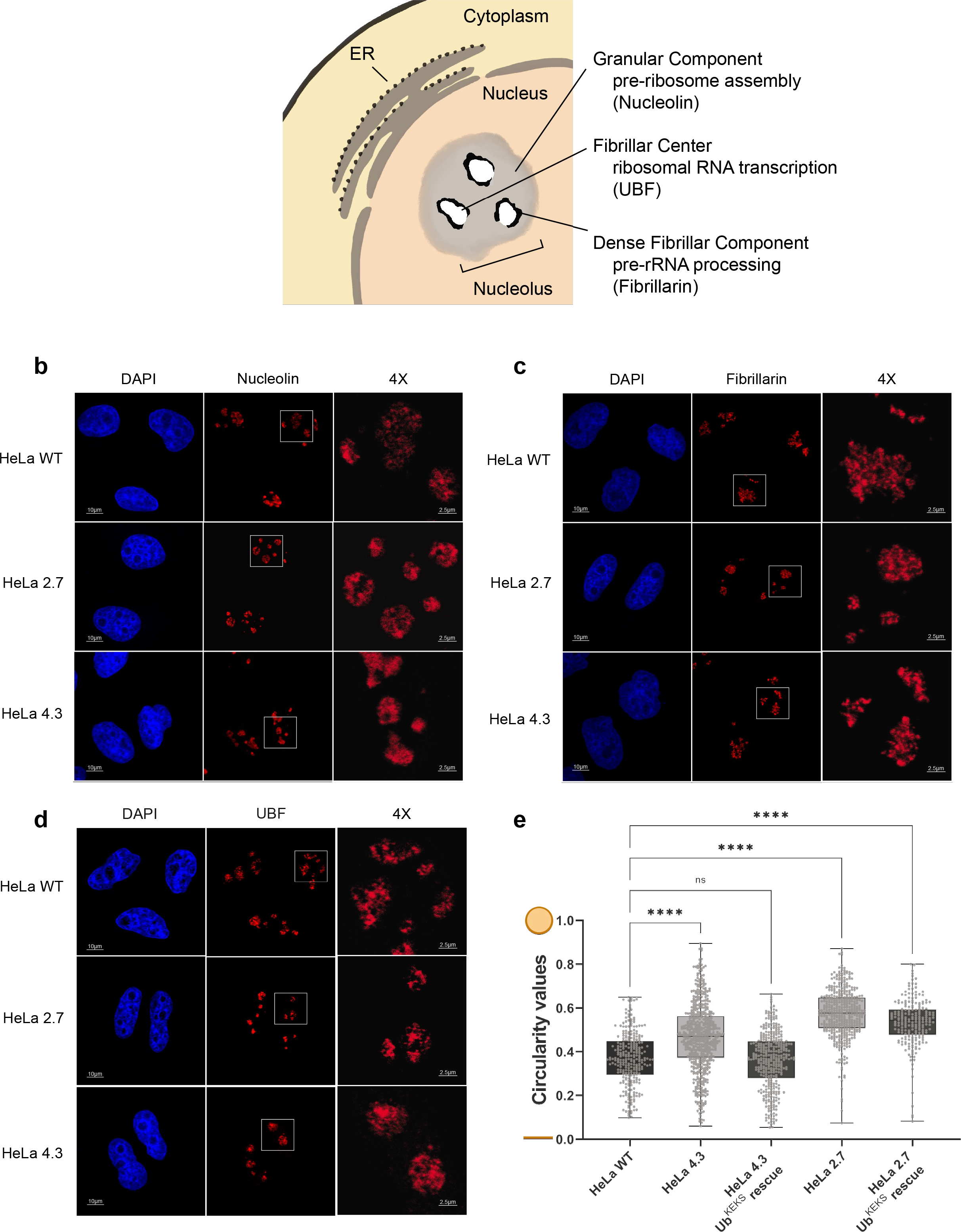
Nucleoli morphology changes upon the loss of Ub^KEKS^. **a.** The nucleolus is a membrane-less organelle composed of 3 distinct sub-compartments: (from the outside to the inner center) the Granular Component, the Dense Fibrillar Component and the Fibrillar Center. The function in ribosomal RNA processing stages and protein markers of each substructure is indicated. **b. c. and d.** Hela WT and Ub^KEKS^ knockout cells (clones 4.3 and 2.7) were labelled for immunofluorescence microscopy with nucleolin, fibrillarin, UBF antibodies and nuclei were stained with DAPI (n=3 independent experiments). The four-times enlarged region (right panel) is delimited by a white box. The scale bar for each field of view is indicated. **e.** Nucleoli circularity measurements were performed in Fiji through object identification using nucleolin. For rescue experiments, transfected cells with Ub^KEKS^ were selected using HA staining. Circularity values are represented on a 0-1 scale where 0 corresponds to a line and 1 to a perfect circle. Data are represented in box plots where the center line shows the medians; box limits indicate the 25th and 75th percentiles as determined by Graph Pad Prism software. Whiskers extend to minimum and maximum values and individual datapoints are superimposed on the box plot. Statistical analysis was performed using one-way ANOVA (F(4,1864)= 186,2) and Dunnett’s multiple comparison post hoc test. “****” indicates an adjusted p-value <0.001. Adjusted p-value for comparison of each knockout clone with their rescue are respectively equal to 0.0029 for HeLa 2.7 clones and inferior to 0.001 for HeLa 4.3 clones. (n= 800 nucleoli, examined over four independent experiments).

### Disturbed nucleoli structure in Ub^KEKS^ knockout cells

HeLa cells lacking the Ub^KEKS^ protein were previously shown to have LMNA foci accumulation in the nucleoli, as well as an increase in the size of the nucleoli, suggesting a potential function for Ub^KEKS^ within this membraneless nuclear structure [14]. To further investigate the difference in nucleolar morphology between wild type and Ub^KEKS^ knockout cells, immunofluorescence analysis was performed using three different nucleolar proteins specific to each sub-compartments of the nucleolus (Fig 2a-d). From the edges to the center, Nucleolin was used for detecting the Granular Component, fibrillarin for the Dense Fibrillar Component and UBF for the Fibrillar Center. While there was no difference in the localization of either of them, the labelling showed that the shape of the nucleoli in the Ub^KEKS^ knockout cells was more rounded with a clearly defined edge. To further study this observation, the circularity of the nucleoli was measured and quantified (Fig 2e). Knockout cells showed significantly more round shaped nucleoli compared to wild type cells, and the reintroduction of Ub^KEKS^ protein through transient transfection rescued or reduced the phenotype in both knockout cell lines (clone 4.3 and clone 2.7 respectively). The morphology of the nucleoli was further investigated using transmission electron microscopy (Fig 3a). Interestingly both Ub^KEKS^ knockout clones showed a disorganized dense fibrillar component (DFC) and fibrillar center (FC) sub-compartments. While in the wild type cells DFC formed a circular structure around the FC, cells lacking Ub^KEKS^ had misshapen structures either with distorted forms or less DFC surrounding the FC region. To address if this structural disturbance has an effect on ribosomal RNA production quantitative PCR (qPCR) was performed for the 45S precursor as well as the 18S and 28S ribosomal RNAs transcribed by RNA Pol I. As a control for RNA Pol II transcription a long non-coding RNA *MALAT1* levels have also been measured (Fig 3b-c). None of the ribosomal RNAs showed difference in transcription in the Ub^KEKS^ knock out cells, suggesting that ribosomal RNA transcription or processing is not affected by the disturbed nucleolar morphology observed.

**Figure 3:**
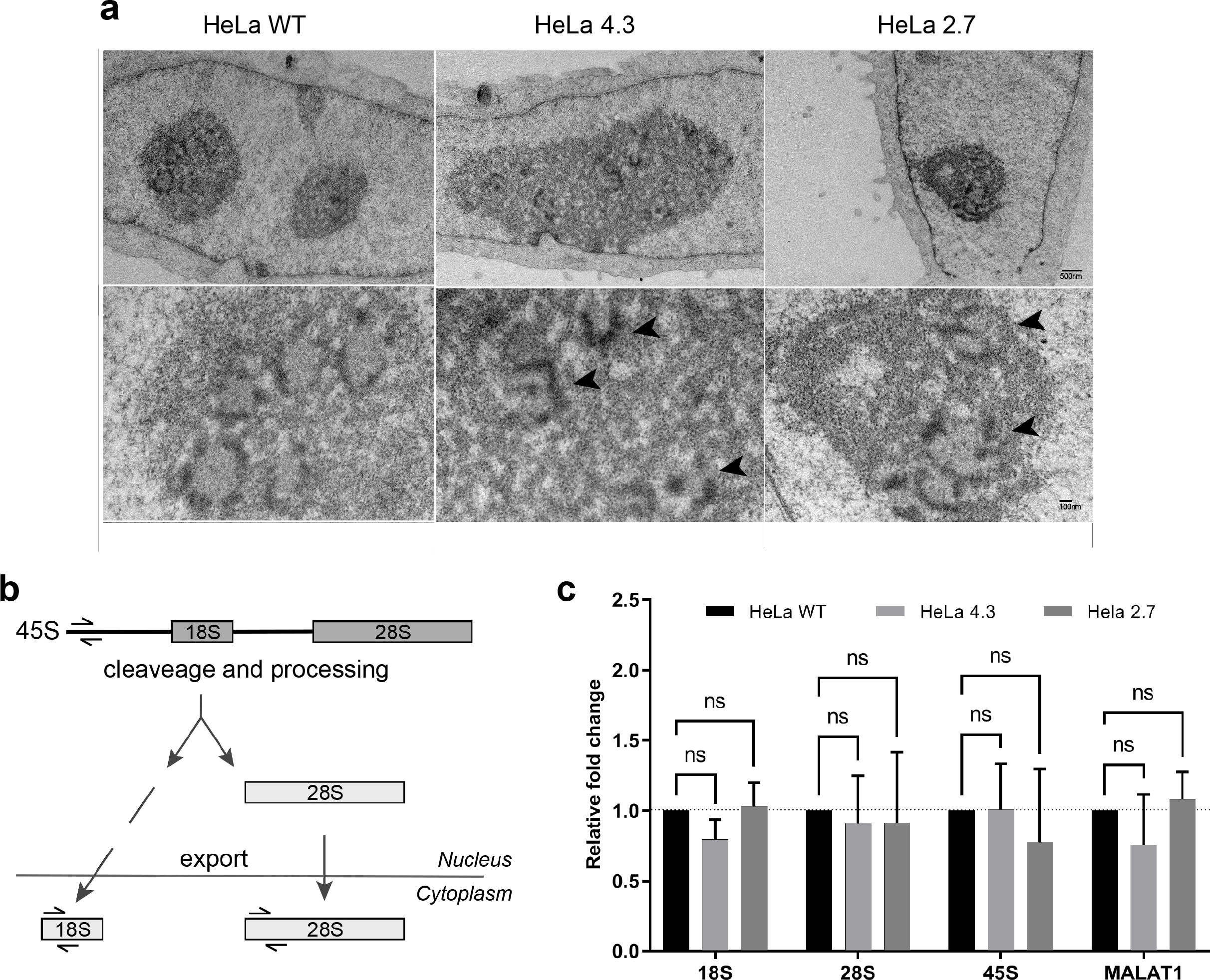
Disrupted nucleoli organization in Ub^KEKS^ knockout cells does not impact ribosomal maturation. **a.** Hela WT and Ub^KEKS^ knockout cells (clones 4.3 and 2.7) were contrasted for electron microscopy with uranyl acetate (n=3 independent experiments). Arrowheads indicate disturbed DFC and FC structures. Scale bars correspond to 500nm (top panels) and 100nm (bottom panels). **b.** Simplified schematic representation of mammalian ribosomal RNA processing. Half arrowheads indicating the qPCR primers used for the amplification of the precursor 45S and mature ribosomal RNAs. **c.** Quantitative measurement of 18S, 28S and 45S pre-RNAs were performed using quantitative PCR (qPCR) analysis. *MALAT1* was used as a control for RNA Pol II transcription. Measurements were performed using LightCycler 96 Instrument and Software. Data were normalised to housekeeping genes and relative expression (compared to HeLa WT samples) were calculated. Data are presented as mean values with ±SD. Statistical analysis were performed using one-way ANOVA (F(2, 6)= 3.024; F(2, 6)= 0.06343; F(2, 6) = 0.4213; F(2, 6) = 1.560) and Dunnett’s multiple comparison post hoc test (n=3 independent experiments).

### Nucleoli morphology differences are associated with a change in proteins composition

Nucleoli from both wild type and Ub^KEKS^ knockout cells were successfully isolated using different sucrose cushions and successive centrifugations (Fig 4a)[42]. Nucleoli isolated from Ub^KEKS^ knockout cells display a significantly larger nucleoli compared to wild type HeLa cells (Fig 4a-4b). It also worth mentioning that the difference of nucleoli’s circularity shown previously is clearly visible in the entire cell pictures (Fig 2e and Fig 4a). DIA mass spectrometry was used once again to identify the differences in protein composition which are responsible for enlarged nucleoli in Ub^KEKS^ knockout cells. Similar to the total cell extracts analysis, a new spectral library was generated from nucleoli isolated from wild type HeLa cells (Fig 4a-Top right picture, Supp Fig 3a). The resulting library displays an enrichment of nucleolar proteins (Supp Fig 3b). Out of the 250 most abundant proteins, 53.2% are known to be located in the nucleolus and 22.40% to the nucleoplasm by current literature. Functional enrichment of those 250 proteins also highlights cellular pathways characteristic of nucleoli, mainly related to ribosome biogenesis, ribonucleoprotein complex biogenesis, non-coding RNA processing or ribosomal RNA maturation (Supp Fig 3c). Correlation coefficients were higher than 0.9 within each cell line, indicating that the isolation of nucleoli and the DIA mass spectrometry analysis were highly reproducible. (Supp Fig 4a). Also, the protein composition of the nucleoli of each knockout clone can clearly be differentiated from wild-type nucleoli as shown by PCA analysis (Supp Fig 4b). Although the absence of Ub^KEKS^ leads to changes in protein composition, it does not result in an imbalance in the overall number of proteins in this organelle (Fig 4c). Among significantly modulated proteins, 7 were down-regulated and 11 were up-regulated in both HeLa 2.7 and HeLa 4.3 (Fig 4d – Supp Fig 4c). Interestingly, 2 additional proteins were found significantly modulated in both Ub^KEKS^ knockout clones but showed opposite trends: ERCCL6 and NCK1 were upregulated in one of the clones while being downregulated in the other (Supp Fig 4c). For protein-protein interactions and functional enrichment analyses, all 20 proteins commons to HeLa 2.7 and HeLa 4.3 were considered. These proteins do not directly interact between one another (Supp Fig 4d) but seem to play a role in apoptosis regulation (Fig 4e).

**Figure 4:**
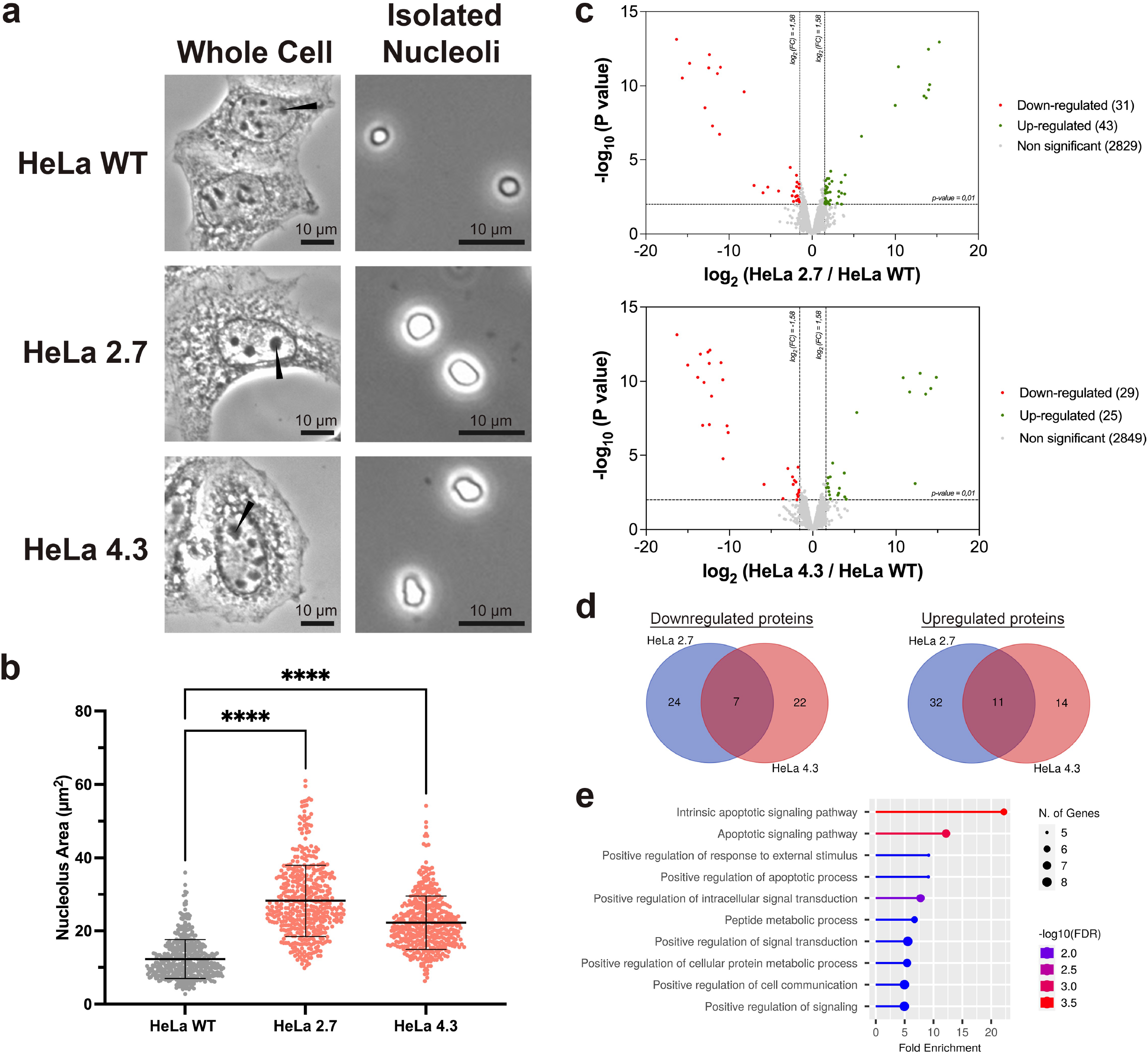
Nucleoli morphology differences are associated with a change of proteins composition. **a.** Phase-contrast microscopy showing exponentially growing HeLa WT, HeLa 2.7 and HeLa 4.3 cells and their isolated nucleoli. Pictures were taken with a 400x magnification for whole cells and with an 800x magnification for the isolated nucleoli. Nucleoli within the cells are indicated by black arrowheads (left panels). Each scale bar represents 10µm (n=3 independent experiments). **b.** Nucleoli area measured with the Fiji software. At least 150 measurements were done for every biological replicate (total number of measurements > 450). Statistical analysis was performed using one-way ANOVA and Dunnett’s multiple comparisons post hoc test (n=3 independent experiments). **c.** Volcano plots showing proteins identified in nucleoli isolated from both wild-type and knockout HeLa cells. Differential abundance analysis was performed using a Limma t-test with slim pi0 calibration. Significantly downregulated proteins are indicated in red (fold change < -3; p-value <0.01) whereas significantly upregulated proteins are shown in green (fold change > 3; p-value < 0.01) (n=3 independent experiments, n=3 independent experiments, each read twice by the mass spectrometer). **d.** Overlaps between significantly downregulated or upregulated proteins in Ub^KEKS^ knockout cell nucleoli. In total, 7 proteins were downregulated and 11 proteins were upregulated in both HeLa 2.7 and HeLa 4.3. Two additional proteins, ERCCL6 and NCK1, were significantly modulated in both Ub^KEKS^ knockout clones. ERCCL6 and NCK1 are however not represented in these Venn diagrams as they were upregulated in one of the clones while being downregulated in the other. The overlapping proteins list was generated using an online Venn diagram tool (Supp Fig. 4c). **e.** Functional enrichment analysis of significantly modulated proteins commons to both HeLa 2.7 and HeLa 4.3. This analysis was done with the online tool ShinyGO (version 0.76) using a False Discovery Rate cut-off of 0,05 combined with the GO biological Process Pathway database. An interaction network of significant proteins was also generated and can be found in Supp Fig. 4d.

### Nucleoli’s protein composition in Ub^KEKS^ knockout cells increases apoptosis

Among the proteins differentially regulated between KO cells and WT cells (Supp Fig 4c), IFI16 and p14ARF (*CDKN2A* gene) were chosen due to their involvement in apoptosis pathway regulation [50][51][52] and used to validate the mass spectrometry data by western blot and immunofluorescence (Fig 5a-b-c). The results confirmed the significant increase in the levels of both proteins in the nucleoli of Ub^KEKS^ knockout clones. Interestingly, such increases are also detectable on a total cell extract level by immunoblotting (Fig 5a) and partially correlate with data from the whole cell proteome mapping: p14ARF was also identified as significantly upregulated by DIA mass spectrometry (“*CDKN2A*” – Supp Fig 2c) whereas IFI16 was labelled as non-significantly modulated. To further investigate functional consequences of these increases in apoptosis-related proteins levels, cellular death was examined in wild type and Ub^KEKS^ knockout cells using Propidium Iodide (PI) and Annexin V labelling by flow cytometry (Fig 5d). A total of four populations were identified due to this double labelling: viable, necrotic, early apoptotic or late apoptotic cells. Necrotic cells percentages were not affected by the presence or absence of Ub^KEKS^. However, Ub^KEKS^ knockout clones showed a significantly higher percentage of apoptotic cells compared to wild type ones. A closer look at all four populations allowed to determine that this apoptotic increase was due to an increase of early apoptotic cells numbers in Ub^KEKS^ knockout clones. Impacts of a higher apoptotic rate on cell proliferation were monitored in real-time by seeding 5 000 wild type or Ub^KEKS^ knockout cells in XCELLigence RTCA E-plates and making regular cellular impedance measurements (Fig 5e). Absence of Ub^KEKS^ strongly decreases cellular growth with notable differences that can be seen as soon as 40 hours after seeding. Pictures of cells colored with Crystal Violet confirmed this proliferation delay due to the lack of Ub^KEKS^ (Pictures on the right panel -Fig 5e). While one week of real-time monitoring allowed wild-type cells to be overconfluent, this time was not even enough for Ub^KEKS^ knockout cells to reach confluency.

**Figure 5:**
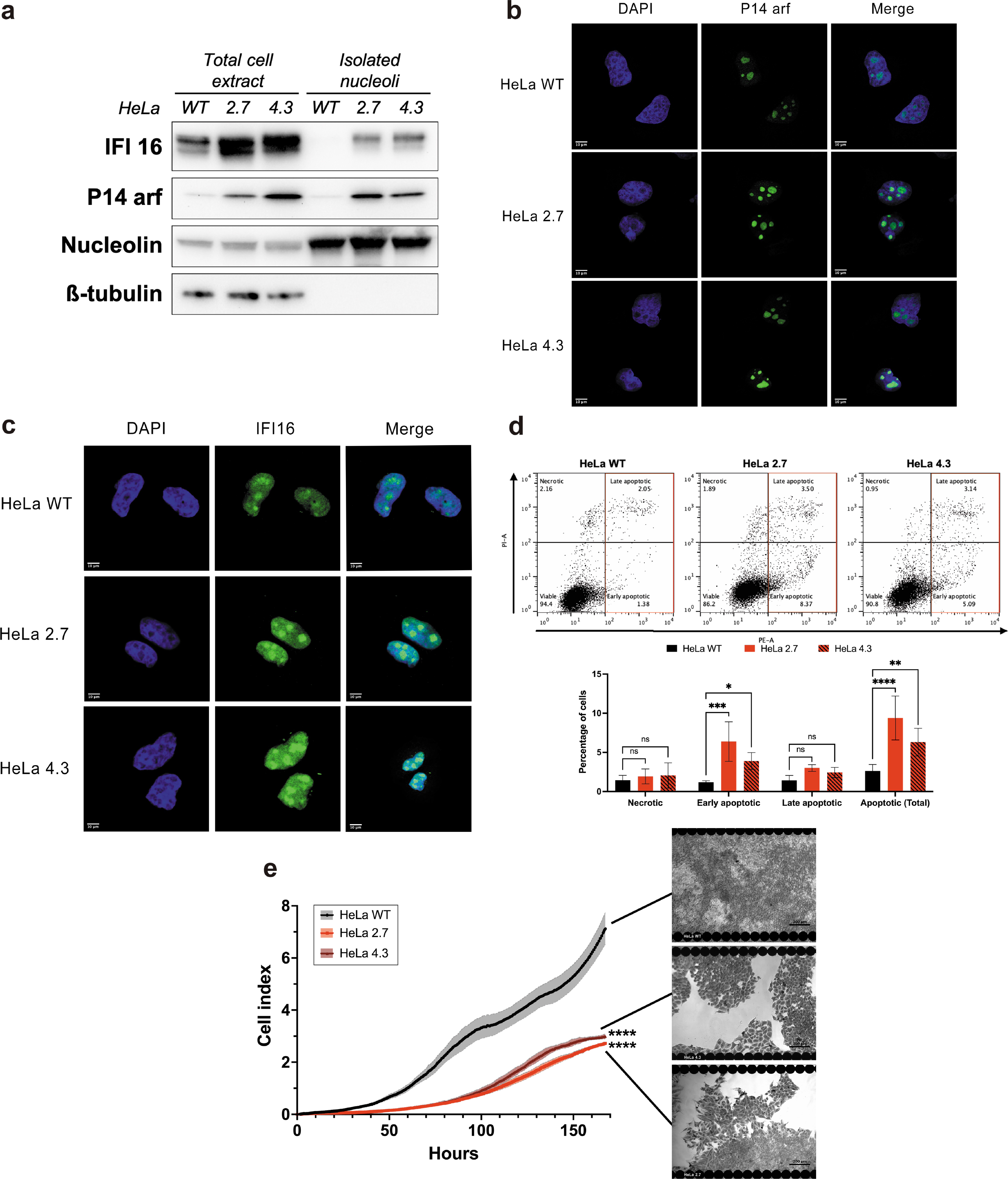
Nucleoli’s protein composition in Ub^KEKS^ knockout cells increases apoptosis. **a.** Total cell extracts and isolated nucleoli from both wild type and knockout HeLa cells were resolved by SDS-PAGE. Significant enrichments of nucleolar proteins which were identified in knockout cells by MS were confirmed with IFI16 and p14ARF antibodies. Membrane was also probed with nucleolin and ß-tubulin antibodies, respectively as loading control and nucleoli extract purity control (n=3 independent experiments). **b. and c.** HeLa WT, HeLa 2.7 and HeLa 4.3 were observed under immunofluorescence microscopy using anti-IFI16 and anti-p14arf antibodies. Cells were stained with DAPI to confirm nuclear and nucleolar location. Scale bar represents 10µm (n=3 independent experiments). **d.** Apoptosis monitoring using Annexin V (coupled to PE fluorophore) and PI labelling. Dot plots present representative data obtained by flow cytometry across all replicates. Gating was performed to identify 4 distinct populations: viable (PI-/PE-), necrotic (PI+/PE-) and early (PI-/PE+) or late (PI+/PE+) apoptotic cells. Gating for total apoptosis (early and late) is indicated in red. Percentage of cells in each population was used for statistical analysis. Significance was determined by two-way ANOVA and Dunnett’s multiple comparisons post hoc test (n=3 independent experiments). **e.** Proliferation curves by cellular impedance. 5000 cells were seeded per well for each type of HeLa cells and cellular impedances were measured every 15 minutes for 1 week. Statistical analysis was done using a two-way ANOVA with Dunnett’s multiple comparisons post hoc test. **** indicates a p-value inferior to 0.0001, representing the cell lines effect on the data. Pictures show confluency reached at the end of the experiment: here, cells were stained with crystal violet for better contrast. Black rounds on the edges of each picture are the electrodes from XCELLigence RTCA E-plates. Scale bar represents 200µm (n=3 independent experiments).

## Discussion

For decades, Ub and Ub-like proteins regulate countless cellular mechanisms by forming complex messages through PTMs [3][11][12][13]. The discovery of Ub variants encoded by what were thought to be pseudogenes adds yet an additional level of regulation to this family of PTMs. Here, we first showed that deletion of Ub^KEKS^ does not result in a significant global change in the whole cellular proteome. This observation supports a different cellular role for Ub^KEKS^ than proteasomal degradation. Meanwhile, at a more precise scale, we also demonstrated that the nucleolus is a cellular compartment particularly impacted by the absence of Ub^KEKS^. Changes in the morphology and internal structure of the nucleoli were observed in Ub^KEKS^ knockout cells, including a bigger and more circular nucleoli and structural disorganization of dense fibrillar component and fibrillar center. Previous studies have already linked nucleolar morphology to environmental stimuli and molecular composition ([16][17][18][20][53]). Here, mass spectrometry analysis revealed that the absence of Ub^KEKS^ results in nucleolar protein composition changes, specifically pointing toward proteins involved in cell cycle regulation and stress response mechanisms. In consequences, we also observed an increase in basal levels of apoptosis and a significant cell proliferation delay in Ub^KEKS^ knockout cells.

DIA mass spectrometry on total cell extracts showed that absence of Ub^KEKS^ does not impact protein stability or degradation. This confirms previous literature work where inhibition of the proteasome by MG132 failed to induce the accumulation of proteins modified by Ub^KEKS^ in cytoplasmic foci, contrarily to proteins modified with Ub [14]. Although proteasomal degradation is the most studied Ub function, many others mechanisms which do not impact protein levels are also regulated by Ub and Ub-like proteins, including cell cycle control, DNA damage response and intracellular protein trafficking [3][13][54][55]. Since the absence of Ub^KEKS^ triggers the accumulation of LMNA in the nucleolus [14] and changes of the nucleolar proteome (this paper), Ub^KEKS^ may act as a regulator for nucleolar protein trafficking.

Nucleolar structures heavily rely on liquid-liquid phase separation principle for its formation [53]. Multiphase liquid immiscibility is the result of biological molecules’ condensation in denser phase, stabilized by intra-macromolecular interactions [56]. Many factors, including intracellular pH and the content of proteins and RNA in the membrane-less compartment, can impact multiphase liquid immiscibility [20][53][57][56][58]. Regarding macromolecules, proteins concentration, protein-protein or protein-RNA interactions, presence of intrinsically disordered regions in protein and PTMs were characterized as key elements to gain correct nucleolar concentric structures [53][57][56][58]. As Ub^KEKS^ appears to be an important PTM for nucleolar organization [14], its depletion results in nucleolar disorganization thus disrupting one of the main criteria of the liquid-liquid phase separation principle. Although nucleolar functions like protein sequestration or DNA repair do not require the correct concentric structure of this cellular compartment, ribosomal biosynthesis is highly dependable on it [16][21][24][53]. Here, we showed by qPCR that ribosomal RNA maturation is not impacted by the absence of Ub^KEKS^ and that only one ribosomal protein paralog (RPL22L1) was significantly modulated by Ub^KEKS^ as shown by DIA mass spectrometry. This protein, however, is not involved in ribosomal biogenesis either [59]. As mentioned previously, the hypothesis that Ub^KEKS^ could be a nucleolar protein trafficking regulator is quite appealing. Those new insights favor this assumption as impairment of proteins exchange between nucleolus and nucleoplasm leads to changes in nucleolar composition, themselves leading to a disruption of the multiphase liquid immiscibility and disorganized nucleoli. Since the absence of Ub^KEKS^ has no impact on ribosomal biogenesis, a wider approach by DIA mass spectrometry was chosen to map the consequences on nucleolar proteins trafficking.

DIA mass spectrometry is a technique that fully developed over the last decade [60]. In contrast to the more common data dependent acquisition (DDA) mass spectrometry which only fragments and analyzes peptides with high intensity, DIA mass spectrometry process and investigates all peptides in a given sample [60][61][62][63]. This allows the generation of more accurate and reproducible data [61]. As DIA can analyse a complete mixture of peptides, it provides a precise, complete and unbiased map of the nucleolar proteome in the presence or absence of Ub^KEKS^. Moreover, DIA uses a peptide library that is generated from a collection of samples, which can either be very comprehensive, but also very specific to a subset of proteins of interest. Here, several proteins whose nucleolar localizations are impacted by Ub^KEKS^ were identified using such a library that was specifically generated from fractionated nucleoli. Functional enrichment of significantly modulated proteins in absence of Ub^KEKS^ pointed toward an activation of apoptosis. The levels of P14arf and IFI16, two proteins involved in the regulation of P53-dependent apoptosis [50][51][52], were higher in the nucleoli of Ub^KEKS^ knockout cells . IFI16 can directly bind P53 and promote phosphorylation on its serine 15 or serine 392, leading to P53 stabilization and apoptosis [51][64][65]. P14arf is an important sensor of different types of cellular stresses, and is thus positively regulated in response to different oncogenic signals [66]. P14arf is normally expressed at low levels because of N-terminal ubiquitination and proteasomal degradation, as well as sequestration within the nucleolus by interaction with NPM [67]. P14arf can trigger apoptosis by binding and sequestering the ubiquitin ligases Mdm2 and ARF-BP1 in the nucleolus, thus stabilizing P53 in the nucleoplasm [50][66][68], but also shows tumor-suppressive activities that are independent of p53 through interactions with TIP60, TOPO I and C1QBP [69][70][71]. The nucleolar retention of P14arf is thought to prevent these tumor suppressor activities, but these mechanisms leading to the cellular redistribution of P14arf have not yet been elucidated. Overall, this suggests a p53-independent role in the cellular stress response in the absence of Ub^KEKS^, since HeLa cells used in this study do not express a functional p53. The stabilization of these proteins in the absence of Ub^KEKS^ suggest that this modification is involved, either directly or indirectly, in regulating their expression and activation.

To conclude, this work demonstrates the role of ubiquitin variant Ub^KEKS^ in protein trafficking, more specifically between the nucleoli and nucleoplasm. By modifying its protein content, Ub^KEKS^ is required for maintaining the concentric structure of the nucleolus. Proteins release from or sequestration in nucleoli by Ub^KEKS^ is shown to have cellular-wide impacts such as apoptosis regulation in a p53-independent manner. This marks the first step for establishing Ub^KEKS^ as a new layer of regulation in the ubiquitination system. Yet, the expression among different tissues and the large number of targets of this ubiquitin variant remind us that many others potential functions of Ub^KEKS^ are waiting to be discovered.

## Acknowledgements

J.F. is a recipient a FRQS studentship. Funding was provided from the Canadian Institutes for Health Research, grant number #398925 to F.-M.B. F.M.B. is a FRQS Senior scholar (award number 281824). X.R. is a recipient of a Canada Research Chair in Functional Proteomics and Discovery of Novel Proteins. X.R. and F.M.B. are members of the FRQS-funded “Centre de Recherche du CHUS”.

## Authors contribution

Experimental approaches were designed by J.F., F-M.B. and X.R.. J.F. performed all experiments presented in Fig 1, 4, 5a, 5d, and 5e, as well as the ones available in all supplementary figures. A.M. completed experiments showcased in Fig 2 and 3. G.M. did immunofluorescences displayed in Fig 5b and 5c. D.L. performed all mass spectrometry acquisitions. Original draft for this manuscript was written by J.F. while reviewing and editing was done by J.F, X.R. and F.-M.B.. All authors have read and agreed to the final version of the manuscript.

## Images license

Some images used in this work were created using the SMART medical art platform from Servier (https://smart.servier.com/). Raw images are licensed under a Creative Commons Attribution 3.0 Unported License. Details of this license can be found at https://creativecommons.org/licenses/by/3.0/

## Data availability

The mass spectrometry proteomics data have been deposited to the ProteomeXchange Consortium via the PRIDE [72] partner repository with the dataset identifier PXD040778. Mass spectrometry data for the human cell lines library used in figure 1 is also available under the dataset identifier PXD040784.

---

**Supplementary Figure 1:**
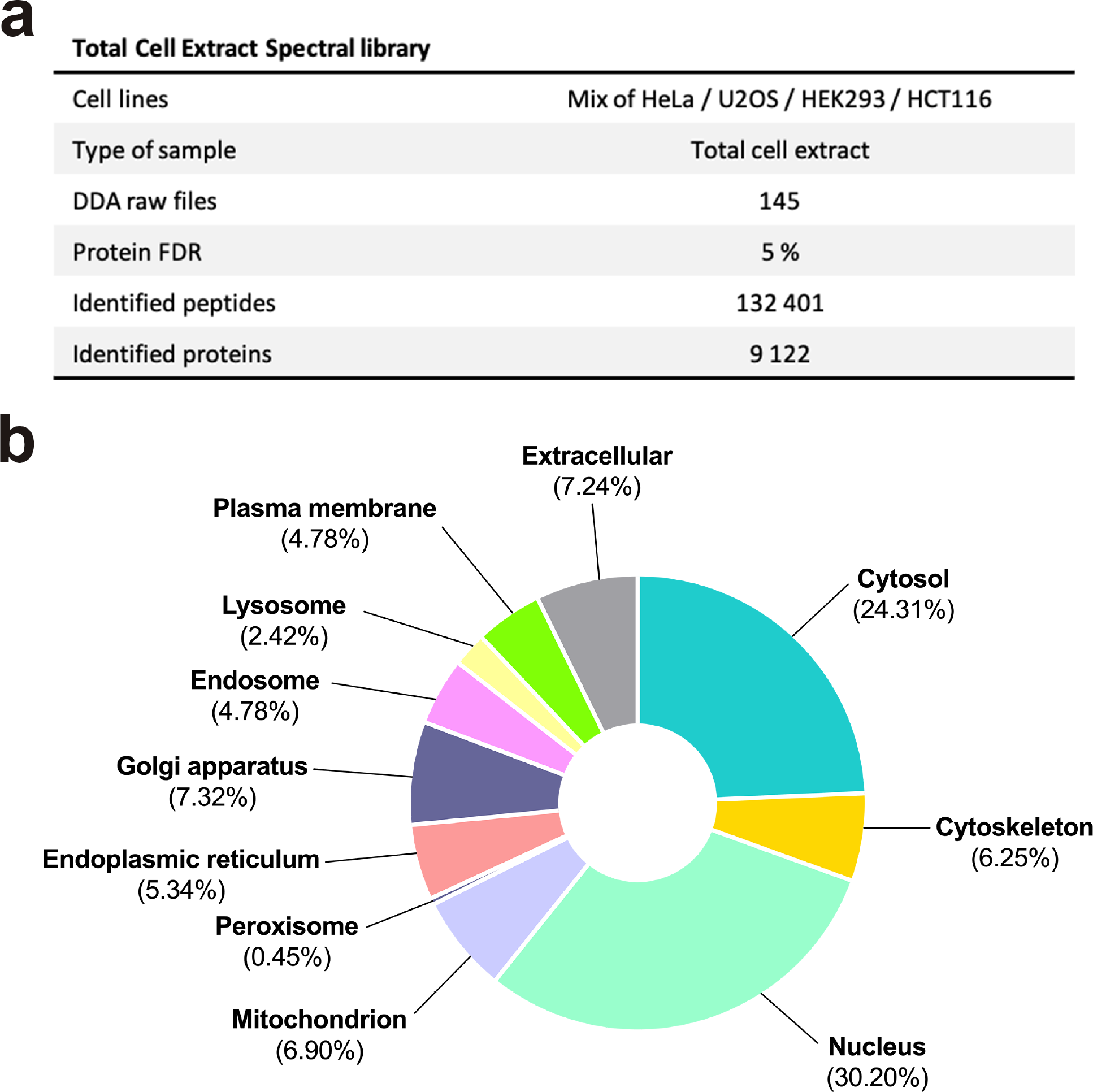
Total cell extracts spectral library. **a.** Main characteristics of the spectral library generated for total cell extracts DIA mass spectrometry. **b.** Representative distribution of proteins identified during spectral library generation. Location was determined using the COMPARTMENTS prediction tool with Cytoscape. Only locations with a false discovery rate inferior to 5% were considered.

**Supplementary Figure 2:**
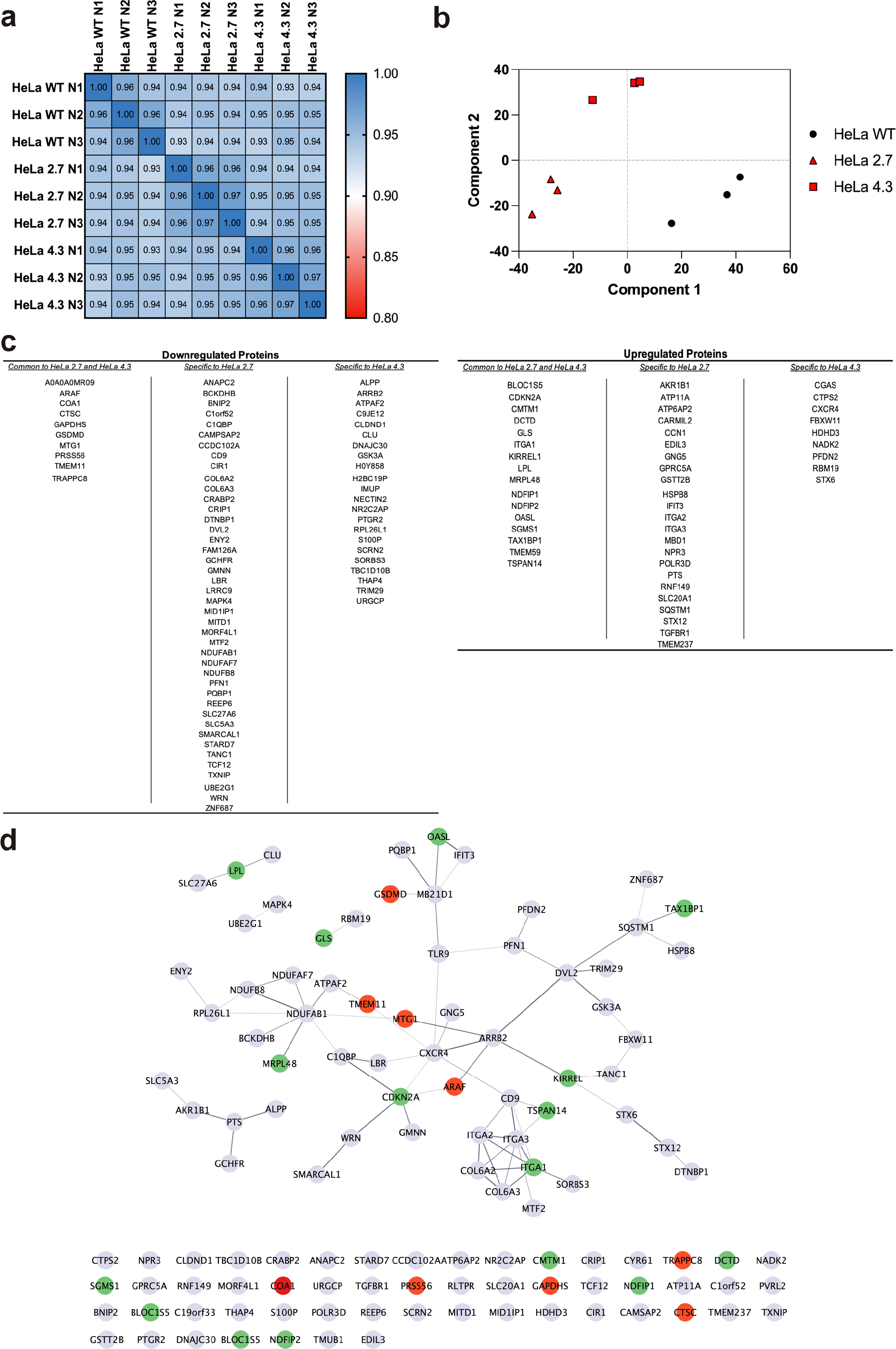
Complementary data for the DIA run on total cell extracts. **a.** Pearson’s correlation matrix for each replicate of the experiment (n=3 independent experiments, each read twice by the mass spectrometer). **b.** Principal Component Analysis (PCA) using Benjamini-Hoechberg cutoff algorithm (False discovery rate = 5%). **c.** Complete list of proteins significantly modulated in Ub^KEKS^ knockout cells. Overall, 74 proteins are downregulated and 48 proteins are upregulated in HeLa 2.7 and HeLa 4.3. **d.** Interactions network of proteins significantly downregulated or upregulated in Ub^KEKS^ knockout cells. Downregulated and upregulated proteins common to both HeLa 2.7 and HeLa 4.3 are coloured in red and green respectively. A functional enrichment analysis was also performed with the online tool ShinyGO (version 0.76) using a False Discovery Rate cut-off of 0,05 combined with the GO biological Process Pathway database. No significant functional enrichment was retrieved using these parameters (not shown).

**Supplementary Figure 3:**
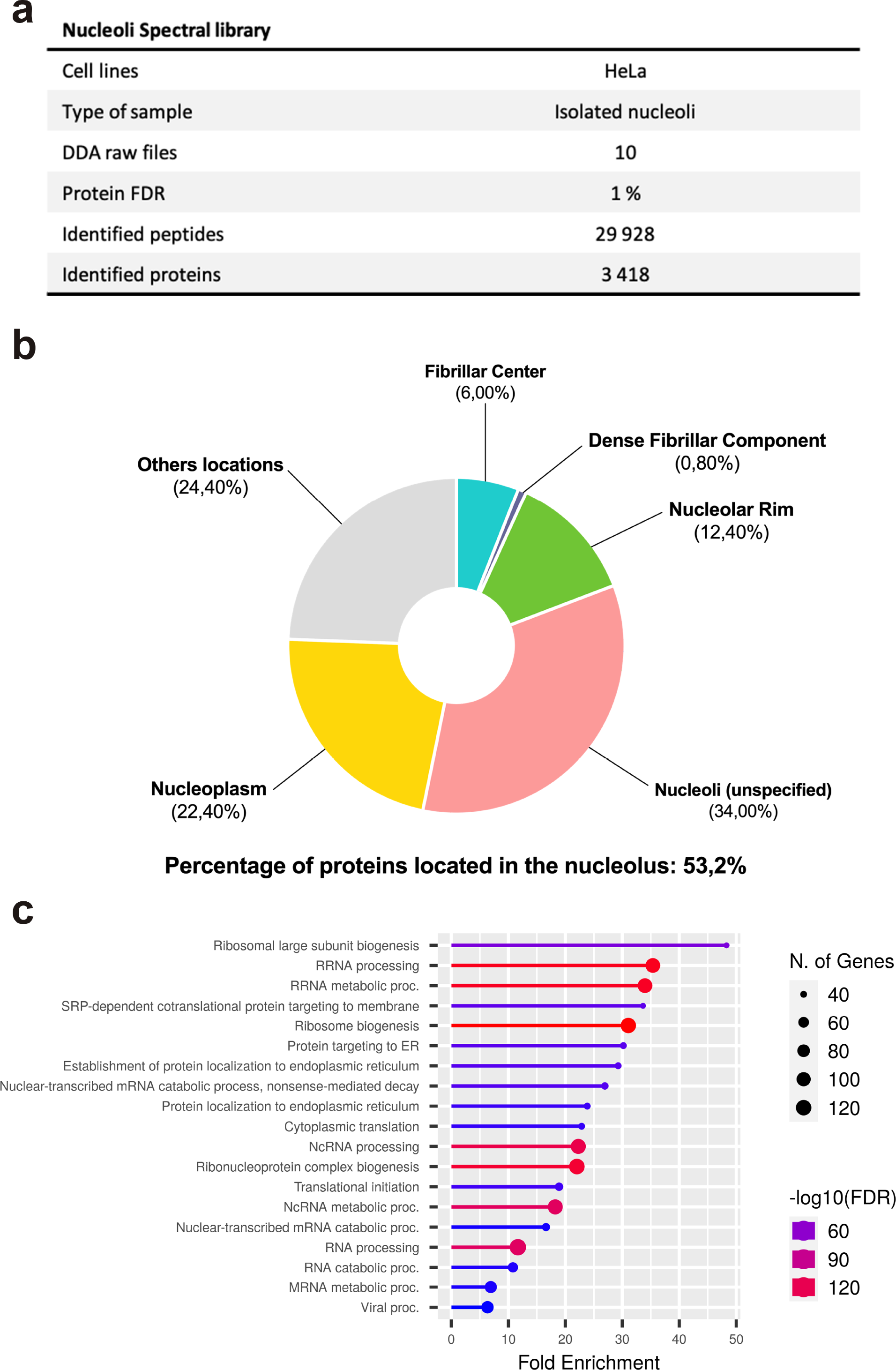
Nucleolar spectral library. **a.** Main characteristics of the spectral library generated for DIA mass spectrometry of isolated nucleoli. **b.** Representative distribution of the 250 most abundant proteins identified during spectral library generation. Location was determined using the COMPARTMENTS prediction tool with Cytoscape. Only locations with a false discovery rate below 5% were considered. **c.** Functional enrichment for the 250 most abundant proteins of the nucleolar spectral library. Analysis was performed via the online tool ShinyGO (version 0.76) with a False Discovery Rate cut-off of 0,05 using the GO biological Process Pathway database.

**Supplementary Figure 4:**
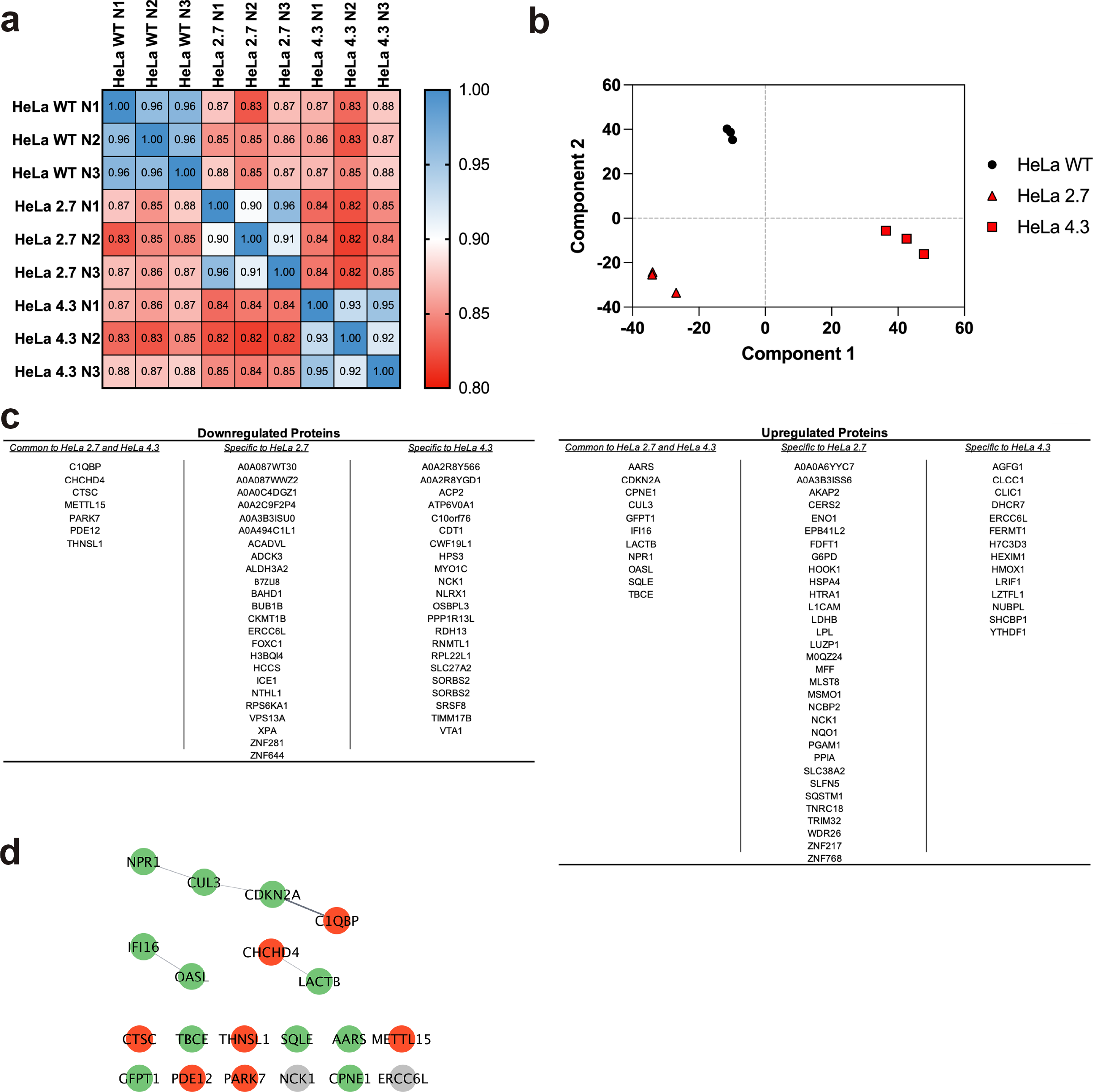
Complementary data for the DIA run on isolated nucleoli. **a.** Pearson’s correlation matrix for each replicate of the experiment (n=3 independent experiments, each read twice by the mass spectrometer). **b.** Principal Component Analysis (PCA) using Benjamini-Hoechberg cutoff algorithm (False discovery rate = 5%). **c.** Complete list of proteins significantly modulated in Ub^KEKS^ knockout cells. Overall, 53 proteins are downregulated and 57 proteins are upregulated in HeLa 2.7 and HeLa 4.3. **d.** Interactions network of proteins significantly downregulated or upregulated in both Ub^KEKS^ knockout cells. Downregulated and upregulated proteins common to both HeLa 2.7 and HeLa 4.3 are coloured in red and green respectively. Proteins in grey were significantly upregulated in HeLa 2.7 but significantly downregulated in HeLa 4.3.

